# Zinc^2+^ ion inhibits SARS-CoV-2 main protease and viral replication *in vitro*

**DOI:** 10.1101/2021.06.15.448551

**Authors:** Love Panchariya, Wajahat Ali Khan, Shobhan Kuila, Kirtishila Sonkar, Sibasis Sahoo, Archita Ghoshal, Ankit Kumar, Dileep Kumar Verma, Abdul Hasan, Shubhashis Das, Jitendra K Thakur, Rajkumar Halder, Sujatha Sunil, Arulandu Arockiasamy

## Abstract

Zinc deficiency is linked to poor prognosis in COVID-19 patients while clinical trials with Zinc demonstrate better clinical outcome. The molecular target and mechanistic details of anti-coronaviral activity of Zinc remain obscure. We show that ionic Zinc not only inhibits SARS-CoV-2 main protease (Mpro) with nanomolar affinity, but also viral replication. We present the first crystal structure of Mpro-Zn^2+^ complex at 1.9 Å and provide the structural basis of viral replication inhibition. We show that Zn^2+^ coordinates with the catalytic dyad at the enzyme active site along with two previously unknown water molecules in a tetrahedral geometry to form a stable inhibited Mpro-Zn^2+^ complex. Further, natural ionophore quercetin increases the anti-viral potency of Zn^2+^. As the catalytic dyad is highly conserved across SARS-CoV, MERS-CoV and all variants of SARS-CoV-2, Zn^2+^ mediated inhibition of Mpro may have wider implications.

## Main Text

COVID-19 pandemic caused by SARS-CoV-2 is a major clinical challenge ^1 2 3^. Lower serum Zinc concentration at the time of admission of COVID-19 patients correlates with severe clinical presentations; an extended duration to recovery, higher morbidity, and a higher mortality in elderly ^4 5^ . However, clinical trials with Zinc and ionophore show positive clinical outcome with a decreased rate of mortality, and transfer to hospice ^6 7^ ^8^

Zinc plays several key roles in biological systems viz. structural, catalytic, regulatory and signalling events ^9 10 11^ . Further, Zinc exhibits anti-viral properties ^12^, including against SARS-CoV. SARS-CoV Main protease (Mpro) ^13^ and RNA dependent RNA polymerase (RDRP) ^14^ are potential key molecular targets of Zinc. However, the structure of SARS-CoV-2 RDRP ^15^ suggests a structural role for Zinc rather than an inhibitory one. Notably, detailed kinetics and mechanism of ionic Zinc targeting SARS-CoV-2 Mpro is lacking.

We first studied one on one binding kinetics of Zinc acetate with purified SARS-CoV-2 Mpro using Surface Plasmon Resonance (SPR). Zinc binds to SARS-CoV-2 Mpro with an association rate constant (ka) of 8,930±30 M^-1^s^-1^ and the dissociation rate constant (kd) of 0.01755±10 s^-1^, and an equilibrium dissociation constant (KD) of 1.965E-06 M. (**Fig. 1a**) The half-life (t1/2=ln [0.5]/kd) of Mpro-Zn^2+^ complex is ∼40s. We then assessed the inhibitory effects of Zn^2+^ binding on the proteolytic activity of SARS-COV-2 Mpro in the presence of Zinc acetate. We obtained an IC_50_ value of 325.1 ± 5.1 nM with complete inhibition at 6.25 µM and above (**Fig. 1b**). We also tested Zinc glycinate and Zinc gluconate complexes, which are available as Zinc supplements in the market and are also investigated in COVID-19 clinical trials^16^, and obtained IC_50_ values of 279.35±17.95 nM and 405.25±0.45 nM, respectively (**Supplementary Figure 1a and 1b**). Reversibility of Zn^2+^-mediated inhibition was tested by first inhibiting the enzyme with 500 nM Zinc acetate, and then initiating the reaction with a substrate peptide, followed by addition of EDTA to regain the enzyme activity by chelating Zn^2+^ ions. We find that Zinc inhibition is completely reversible by EDTA (**Supplementary Figure 1c**), suggesting that inhibition by the metal ion is not because of oxidation of catalytic cysteine (Cys145).

**Figure 1.**
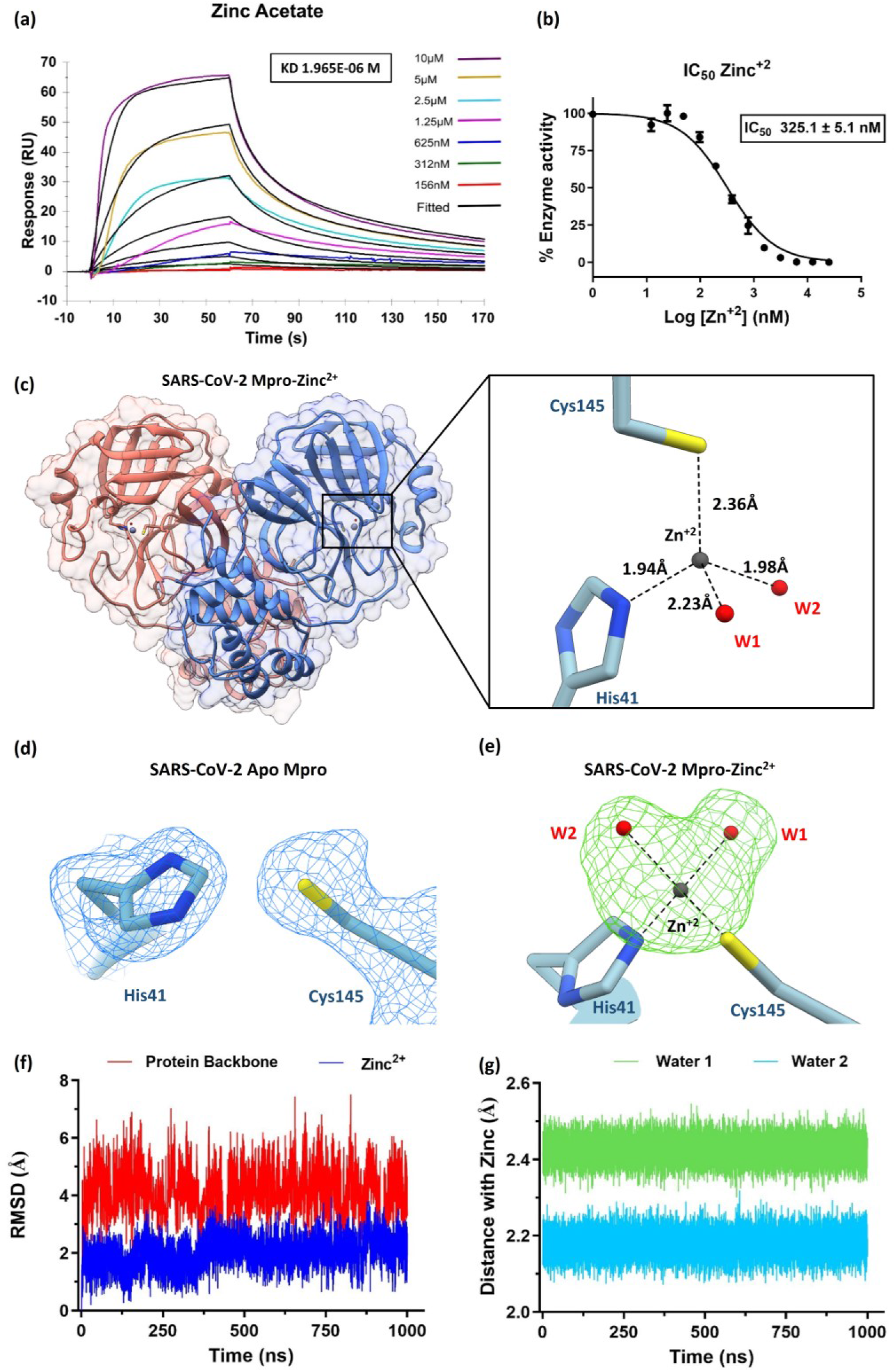
Zn^2+^ binds at the active site and inhibits SARS-CoV-2 Mpro enzyme activity: **(a)** Interaction kinetics of Zn^2+^ with immobilized Mpro using surface plasmon resonance (SPR) shows affinity (KD) of 1.96 µM. Coloured lines indicate various concentrations of Zinc acetate. **(b)** IC_50_ and concentration dependent inhibition of Mpro by Zn^2+^ ion. **(c)** Complex crystals structure of Mpro dimer with Zinc (grey sphere) bound at the active site of both protomers. On the right, catalytic dyad Cys145 and His41 of Mpro is shown with bound Zinc in tetrahedral coordination geometry. **(d)** Electron density map (2Fo-Fc) contoured at 1 σ showing the catalytic dyad in Apo-Mpro **(e)** Omit difference map (Fo-Fc contoured at 3 σ) shows unambiguous density (green) for Zn^2+^ (grey) and two metal-ion coordinating structural water molecules (red). **(f)** 1µs MD simulation Mpro-Zn^2+^ complex shows low RMSD of Zn^2+^ (blue) and protein backbone (red) indicating stability of inhibited state **(Supplementary video) (g)** Distance plot shows less fluctuations in inter-atomic distances between Zn^2+^ and two coordinating water molecules during the course of simulation, representing stable metal ion-water interactions in the inhibitory role of Zn^2+^.

To further understand the structural basis of SARS-CoV-2 Mpro inhibition by Zn^2+^ ion, we solved the crystal structure of the bound complex at 1.9 Å (**Supplementary Table 2**). The asymmetric unit contains a dimer of Mpro in space group P2_1_2_1_2_1_ (**Fig. 1c**). An unambiguous electron density for Zn^2+^ (**Fig. 1d, e**) shows that the metal ion is coordinated by the catalytic dyad His41 and Cys145, which is absent in the control datasets collected for apo-enzyme crystals grown in the same condition. Zn^2+^-bound complex shows a tetrahedral coordination geometry at the Mpro active site by coordinating with two water molecules that are absent in the apo-enzyme structure (**Fig. 1c**). Distortion in the tetrahedral geometry observed is attributed to the presence of heterogeneous atoms; sulphur (Cys145-SG) and nitrogen (His41-NE2) in the inhibited complex. A 180° flip of the imidazole ring of His41 brings NE2 closer to Zn^2+^ with an interatomic distance of 1.94 Å to form a coordinate bond. The interatomic distance between catalytic Cys145 and Zn^2+^ is 2.36 Å consistent with observed bound complexes. Two structural water molecules W1 and W2 (PDB: 7DK1; HETATM 5028 and 5031, respectively) coordinate Zn^2+^ at an inter-atomic distance of 2.23 Å and 1.98 Å, respectively, to satisfy the tetrahedral geometry (**Fig. 1c**). The coordination of Zn^2+^ with the catalytic dyad is expected to prevent a nucleophilic attack on the carbonyl moiety of the amide bond in polyprotein substrate. We hypothesize that the two strongly coordinated water molecules impart stability to the Zn^2+^ inhibited complex

To gain deeper insights into the stability of Mpro-Zn^2+^ complex, we simulated the complex for 1 µs at 300K, keeping the coordinating waters, W1 and W2. During the simulation, Zn^2+^ remains strongly bound to the active site via metal coordination bonds with His41 (NE2) and Cys145 (SG) with an interatomic distance of 1.951±0.031 and 2.518±0.031 Å, respectively, throughout the 1μs time frame. The mobility of Zn^2+^ ion is restricted with a mean RMSD of 0.920 Å (**Fig. 1f**) in accordance with the dynamics of side chains of coordinating catalytic dyad. Visualization of MD simulation trajectory shows (**Supplementary Movie**) that coordinating water molecules W1 and W2 remain bound to Zn^2+^ throughout the simulation, and help maintain the tetrahedral geometry observed in complex crystal structure (**Fig. 1g**)

We further assessed the inhibitory potential of Zinc acetate, Zinc glycinate and Zinc gluconate against SARS-COV-2. Infected Vero E6 cells were treated with all the three Zinc salts at their respective maximum non-toxic concentrations (MNTD) for 48 hours. The MNTDs used for the three compounds were 100 µM for Zinc acetate and Zinc gluconate and 70 µM for Zinc glycinate. We observed that Zinc acetate treatment resulted in more than 50% reduction of viral titre, as compared to the untreated infected cells (**Fig. 2a**). Based on these results, we determined the IC_50_ of Zinc acetate to be 3.227 µM (**Fig 2b**). Surprisingly, Zinc glycinate and Zinc gluconate failed to inhibit viral replication in standard antiviral assays at non-toxic concentrations albeit showing effective enzyme inhibition *in vitro*. Quercetin, a natural Zinc ionophore, increases the bioavailability of Zinc in treated cells^17^, which prompted us to ask whether an increased bioavailability of Zn^2+^ results in enhanced inhibition of viral replication. To test this, we mixed Zinc acetate with Quercetin at 1:2 molar ratio ^18^ at non-toxic concentrations (**Supplementary Figure 2**) and tested the antiviral activity against SARS-CoV-2. We observed >2-fold viral inhibition in the presence of Quercetin (**Fig. 2c**).

**Figure 2.**
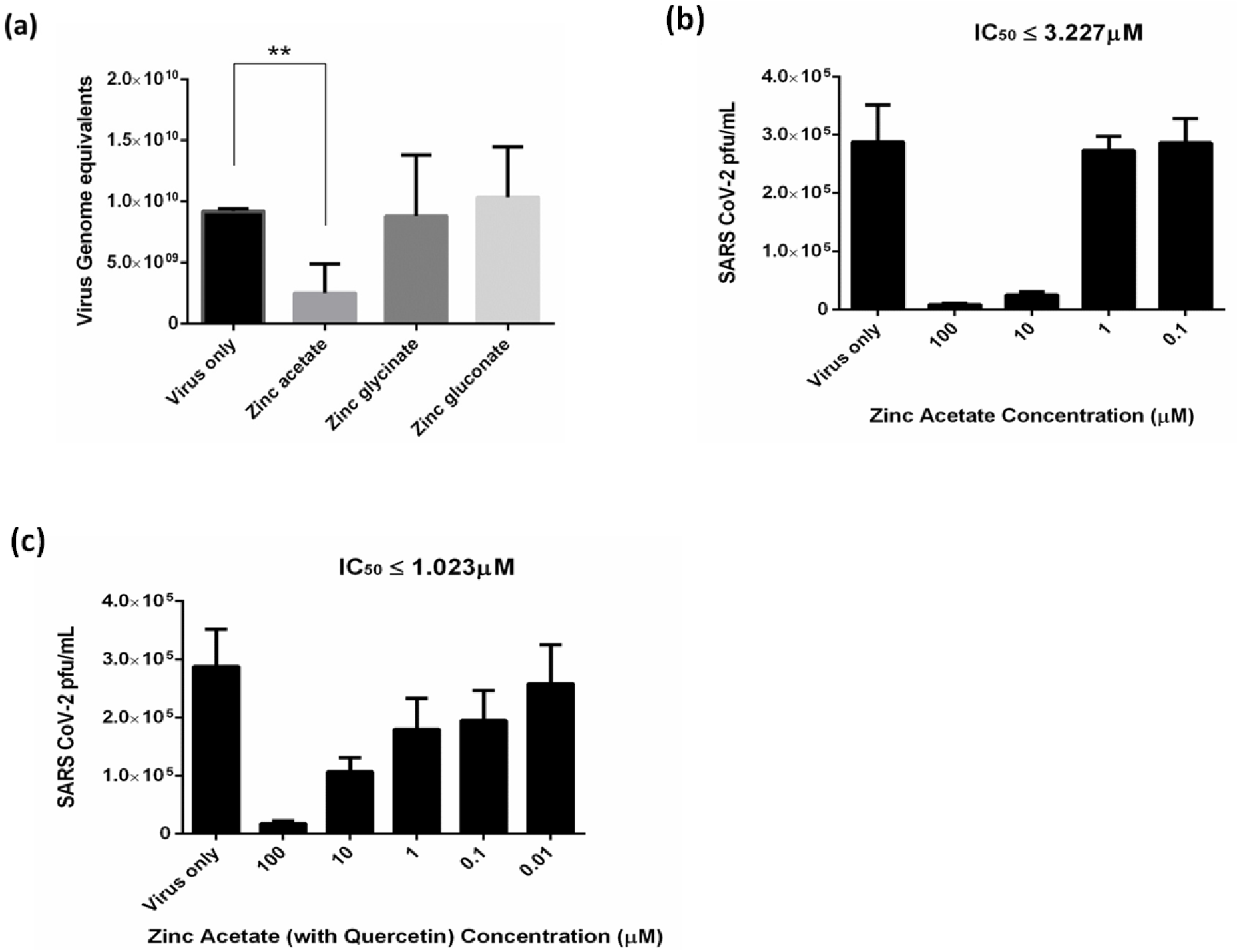
Anti-SARS-CoV-2 activity of Zinc and its complexes with ionophore in infected Vero E6 cells. **a)** Zinc acetate inhibits SARS-CoV-2 replication in Vero E6 cells as determined using qRT-PCR. Treatment with Zinc acetate (100 µM) for 48 h resulted in >50% reduction of viral titre while the Zinc glycinate and Zinc gluconate complexes did not show significant reduction. **(b)** IC_50_ determination using varying concentrations of Zinc acetate for 48 h followed by viral quantification using plaque assay. **(c)** Viral inhibition by Zinc acetate and Quercetin mixture (1:2 M ratio). Mean percentage reduction of SARS-CoV-2 is indicated within the bars. The antiviral experiments were repeated at least twice, and each experiment included at least three replicates. Statistical significance was determined using Student’s t-test (n ≥ 2 biological replicates).

Taken together, our data strongly suggest an inhibitory role for ionic Zinc ^11^, wherein it inhibits SARS-CoV-2 Mpro enzyme activity, supported by complex crystal structure and subsequent inhibition of viral replication *in vitro*. Known crystal structures of Zinc conjugates such as N-ethyl-n-phenyl-dithiocarbamic acid (EPDTC), JMF1600, and Zinc-pyrithione in complex with 3C-like (Mpro) proteases from coronavirus^19^, including SARS-CoV^20^ and SARS-CoV-2^21^, show a similar mode of metal ion coordination with the catalytic dyad (Supplementary Figure 3). However, the Zn^2+^ inhibited SARS-CoV-2 Mpro enzyme structure presented in this study clearly suggests that ionic form of Zinc alone is capable of inhibiting the enzyme, by forming a stable complex at the active site with the help of two water molecules, previously unknown. We further show Zinc complexes; Zinc glycinate and Zinc gluconate failed to show any antiviral effects in our cell culture experiments. Most notably, we show that Zinc ionophore Quercetin aids in inhibition of SARS-CoV-2 replication as it increases the intracellular concentration of Zinc^17^. Our data support the findings that a combination of Zinc salt, which provides ionic Zinc, with ionophores ^6 7 8^, may have a better clinical outcome in COVID-19 therapy. As the Zn^2+^ coordinating catalytic dyad; Cysteine and Histidine is conserved across all coronaviral 3C-like proteases (Mpro), including the variants of SARS-CoV-2, the mode of Zn^2+^ mediated inhibition is expected to be similar. Whether Zn2+ targets Mpro *in vivo* requires further investigation.

## Data accessibility

Mpro-Zinc^2+^ complex coordinates are available at PDB: 7DK1. X-ray raw data is available from Integrated Resource for Reproducibility in Macromolecular Crystallography (IRRMC) repository (https://proteindiffraction.org/).

## Acknowledgement

We thank Prof. Rolf Hilgenfeld, Institute of Biochemistry, University of Lübeck, Lübeck, Germany for providing the expression construct for SARS-CoV-2 Mpro. We thank the beamline staff at the Elettra XRD2 particularly Raghurama P. Hegde and Annie Heroux for beamline support. Access to the XRD2 beamline at the Elettra synchrotron, Trieste was made possible through grant-in-aid from the Department of Science and Technology, India, vide grant number DSTO-1668. The following reagent was deposited by the Centers for Disease Control and Prevention and obtained through BEI Resources, NIAID, NIH: SARS-Related Coronavirus 2, Isolate USA-WA1/2020, NR-52281. We thank Prof. Ramesh Sonti, former Director, NIPGR, New Delhi for access to Biacore T-200. This work was supported by ICGEB core grant and Govt. of India DST-SERB IRHPA grant: IPA/2020/000285.

## Author contribution

LP, WK, SK, KS purified Mpro and performed biochemical and SPR experiments and SD and JKT helped with SPR experiments and data analysis. LP and KS crystallized, collected X-ray data and solved the structure. LP and SS performed MD simulation and analysis. AG performed cytotoxicity assays. AK, DV, AH and S. Sunil performed anti-viral assays and analysed the data. RH provided inputs to biochemical assays. AA coordinated the work and LP and AA wrote the manuscript with inputs from all the authors.

