## Supplementary Figure 1, Supplementary Figure 2, Supplementary Figure 3, Supplementary Table 1, Supplementary Table 2, Materials and Methods for "Zinc^2+^ ion inhibits SARS-CoV-2 main protease and viral replication *in vitro*"

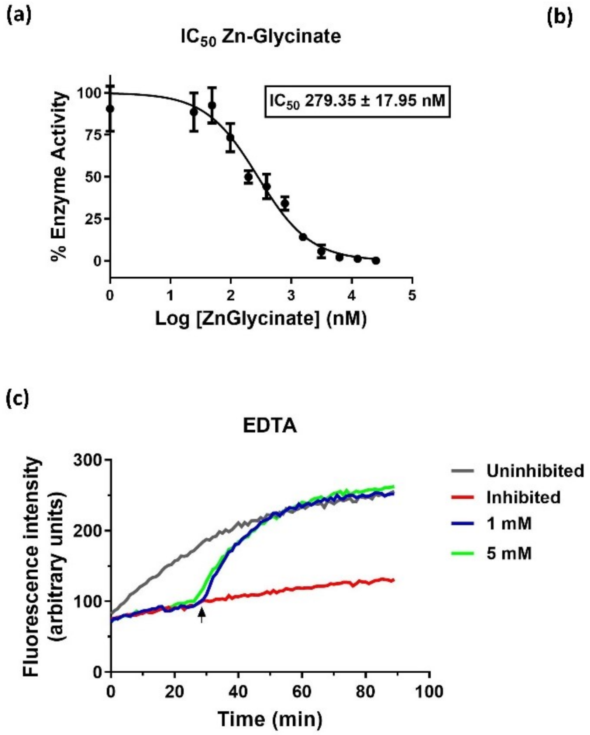

**Supplementary Figure 1. Mpro Inhibition by Zn-complexes.** IC50 and concentration dependent inhibition of Mpro by Zinc-Glycinate (a) and Zinc-Gluconate (b), respectively. (c) Inhibition with 500 nM Zinc acetate was completely reversed by the addition of 1 and 5 mM of EDTA at the $28^{th}$ minute of the ongoing enzymatic reaction.

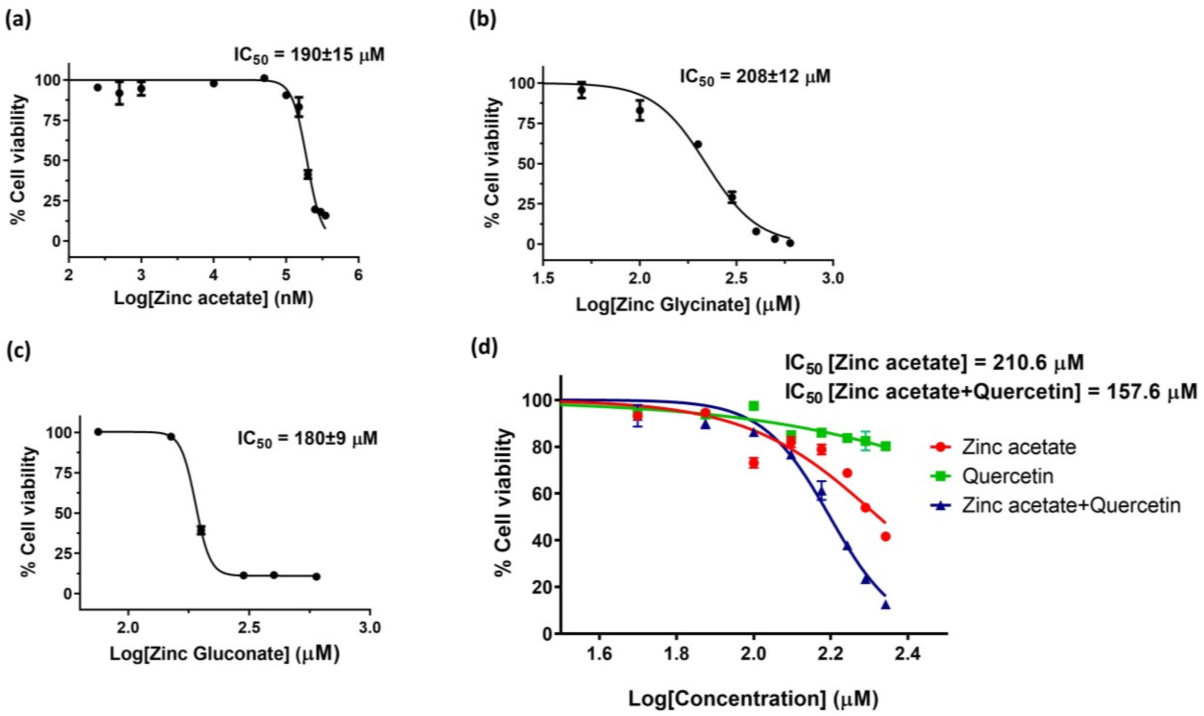

**Supplementary Figure 2. Toxicity determination of Zinc and its complexes in Vero E6 cells.** Non-toxic concentrations were determined by studying the effect of Zinc acetate (a), Zinc glycinate (b), and Zinc gluconate (c) on the proliferation of Vero E6 cells after 48 h post addition, as determined by MTT assays. (d) Non-toxic concentrations for Zinc acetate and quercetin (1:1 molar mixture, blue) is compared with Zinc acetate alone (red). $IC_{50}$ for quercetin alone (green) could not be determined. All experiments are done in biological triplicates.

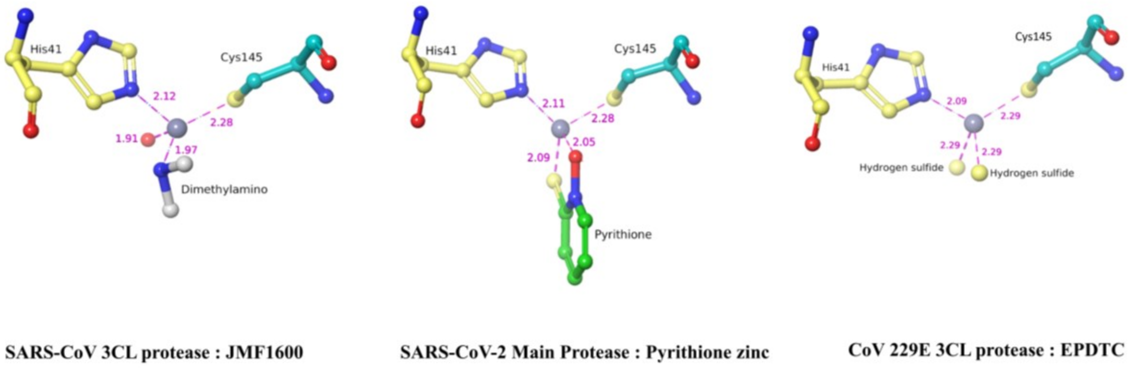

**Supplementary Figure 3: Metal ion coordination of Zinc-complexes bound to coronavirus 3C-like proteases.** Ball and stick model representation of 3CL-pro-Zn complex crystal structures; SARS-CoV-Mpro-JMF1600 (PDB: 2Z9K), SARS-CoV-2-Mpro-Zn-pyrithione (PDB: 7B83) and HCoV-229E-3CLpro-N-ethyl-n-phenyl-dithiocarbamic acid (EPDTC) (PDB: 2ZU2). Zinc is depicted as grey ball. Interatomic distances are represented as dotted lines with bond distance in angstrom (Å).

| Experiment | ka (1/Ms) | kd (1/s) | KD (M) | Rmax (RU) | Chi$^2$ (RU$^2$) | U-value |
|---|---|---|---|---|---|---|
| 1 | 8960 | 0.01745 | 1.95E-06 | 57.32 | 3.39 | 2 |
| 2 | 8899 | 0.01765 | 1.98E-06 | 57.58 | 3.36 | 3 |
| Median | 8,930±30 | 0.01755±10 | 1.965E-06 ±0.15E-07 | 57.45±0.13 | 3.375±0.015 | 2.5±0.5 |

**Supplementary Table 1. Binding kinetics of $Zn^{2+}$ and SARS-CoV-2 Mpro**. Using 1:1 binding model, association [ka (1/Ms)] and dissociation [KD (1/s)] rate constants, and affinity [KD (M)] are shown. The goodness of fit (Chi$^2$ =3.375±0.015) is within the suggested 10% of the Rmax (57.45±0.13). U-values of 2 and 3 reflect 98 and 97% confidence in KD value respectively. The affinity (KD) of $Zn^{2+}$ for Mpro is ~2 μM. The experiment was repeated twice to get the median KD of 1.96 μM.

| | **Mpro-$Zn^{2+}$ (PDB:7DK1)** | **Mpro-Apo (control)** |
|---|---|---|
| **Data collection** | | |
| Space group | P 21 21 21 | P 21 21 21 |
| Cell dimensions | | |
| $a$, $b$, $c$ (Å) | 67.6, 102.2, 102.3 | 67.7, 100.7, 104.0 |
| $\alpha$, $\beta$, $\gamma$ (°) | 90, 90, 90 | 90, 90, 90 |
| $R_{merge}$ | 0.09 | 0.086 |
| Resolution (Å) | 72.32 – 1.90 | 56.79 – 1.81 |
| $I$ / $\sigma I$ | 2.69 (1.9) | 2.44 (1.8) |
| Completeness (%) | 100.0 | 100.0 |
| Redundancy | 12.8 | 12.8 |
| **Refinement** | | |
| Resolution (Å) | 72.32 – 1.90 | 56.79 – 1.81 |
| No. reflections | 56,431 | 66,014 |
| $R_{work}$ / $R_{free}$ | 0.191/ 0.213 | 0.187/0.216 |
| No. atoms | | |
| Protein | 4,582 | 4,649 |
| Ligand/ion | 40 | 38 |
| Water | 423 | 617 |
| $B$-factors | | |
| Protein | 32.4 | 29.4 |
| Zinc ion | 40.75 | - |
| R.m.s. deviations | | |
| Bond lengths (Å) | 0.41 | 0.39 |
| Bond angles (°) | 0.58 | 0.58 |

**Supplementary Table 2. X-ray data processing and refinement statistics**. The data given is for SARS-CoV-2 Apo-Mpro and Mpro-$Zn^{2+}$ complex crystals, both crystallized in the same condition.

### Materials and methods:

**SARS-CoV-2 Mpro purification**: *E. coli* overexpression plasmid pGEX-6p-1 containing SARS-CoV-2 Mpro was a kind gift from Rolf Hilgenfeld, Institute of Biochemistry, University of Lübeck, Lübeck, Germany [1,2]. The procured construct is designed to generate authentic N terminus by auto-proteolytic cleavage via Mpro at the cleavage-site SAVLQ↓SGFRK (arrow represents the cleavage site). Authentic C-terminus was generated by cleaving the C-terminus 6X His-tag at SGVTFQ↓GP by HRV3C protease.

Overexpression and protein purification were performed according to a previous report[2] with some modifications. Expression plasmid was transformed into *E. coli* BL21 (DE3). Transformed cells were inoculated into 200 mL LB media (Luria Bertani Broth, Miller, Himedia) supplemented with ampicillin (100 µg/mL) and grown at 37 °C for 3 h at 100 RPM. The primary culture was used to inoculate 6 L of LB media supplemented with ampicillin and induced with 0.5 mM of isopropyl-D-thiogalactoside (IPTG) after $OD_{600}$ reached 0.8 at 37 °C. 5 h post induction at 37 °C, cells were harvested by centrifugation at 4000 RPM for 20 min at 4 °C and stored at -20 °C until further use. The frozen cell pellet was resuspended in lysis buffer (20 mM Tris, 150 mM NaCl, 10 mM imidazole, 10 µg\ml DNase-I, 100 µg\ml Lysozyme, pH 7.8) and subjected to lysis by sonication on ice, followed by centrifugation at 13000 RPM for 50 min at 4 °C. The supernatant was loaded onto serially connected 2x 5ml HisTrap FF columns (GE) at 0.5 ml/min flow rate, pre-equilibrated with buffer A (20 mM Tris, 150 mM NaCl, 10 mM Imidazole pH 7.8). Non-specifically bound proteins were removed by washing with 5 column volumes (CV) of buffer A. The bound proteins were eluted using buffer B (20 mM Tris, 150 mM NaCl, 500 mM imidazole, pH 7.8) with a linear gradient of 10 to 500 mM imidazole. Fractions containing Mpro were pooled and concentration was estimated using $OD_{280}$ [3] . At this stage, many contaminant proteins were observed. To cleave the C-terminal His-tag, HRV3C[4] protease was mixed with SARS-CoV-2 Mpro in 1:5 ratio (mg/mg) and dialysed into buffer C (20 mM Tris, 150 mM NaCl, 1 mM DTT, pH 7.8) overnight at 4 °C. This was followed by one more round of dialysis for 6 h in buffer A to remove DTT. Dialysed and tag-cleaved protein was passed through serially connected 2x 5ml HisTrap FF columns. Flow through containing enriched Mpro was collected and buffer exchanged with buffer D (20 mM Tris, 1 mM DTT, pH

8.0) using a HiPrep 26/10 desalting column (GE). Desalted protein was loaded onto 5 ml HiTrap Q HP column (GE) pre-equilibrated with buffer D, and eluted using a linear gradient of 0 to 500 mM NaCl in 20 CV of buffer E (20 mM Tris, 1 M NaCl, 1 mM DTT, pH 8.0). Fractions containing pure Mpro were pooled, concentrated and further purified with gel filtration chromatography using pre-equilibrated HiLoad 16/600 Superdex 75 pg column with buffer C at a flow rate of 1 ml/min. Purified protein was concentrated to 27.5 mg/ml, aliquoted and flash frozen in liquid nitrogen and stored at -80 $^{o}$C until further use.

**Surface Plasmon Resonance (SPR):** Experiments were performed using Biacore T200 with control software V2.0 and Evaluation Software V3.1 (GE Life Sciences). All measurements were made at 25 $^{o}$C. Running buffer consisted of HBS-N pH 7.4 (10 mM HEPES, 150 mM NaCl, pH adjusted with NaOH). Purified Mpro was immobilized onto a CM5 chip using amine coupling method according to manufacturer's protocols with 420 s of surface activation with freshly prepared 1:1 mixture of 1-ethyl-3-(-3-dimethylaminopropyl) carbodiimide hydrochloride (EDC) and N-hydroxysuccinimide (NHS) followed by 420 s contact time for the protein over the activated surface at a flow rate of 10 µl/min. The protein (50 µg/ml) was immobilized onto the chip surface in 10 mM acetate buffer pH 4.0 achieving an RU of ~14000. The remaining activated carboxy methyl groups on the surface were blocked by an injection of 1 M-ethanolamine-HCl pH 8.5 for 7 min. An unmodified flow cell surface was used as a reference for each analysis to check for the non-specific binding response to dextran matrix. Running buffer containing varying concentrations of Zinc acetate; 156 nM to 10 µM, were prepared and passed over the immobilized protein at a constant flow rate of 30 µL/min. The interaction (association time 60 s and dissociation time 120 s) between the protein and the analyte resulted in characteristic sensorgrams which were then analysed using Biacore T200 evaluation software; responses generated from unmodified surface were subtracted from the same. The curves (sensorgrams) were fitted using 1:1 model to get the association rate [ka (1/Ms)], dissociation rate [kd (1/s)], and equilibrium dissociation constant [KD (M)] for the interaction. The regeneration was done using 50 mM NaOH, 1 M NaCl solution with 30 s contact time at 30 µL/min flow rate. The experiments were repeated twice to get the median values.

**Mpro enzyme inhibition assay**: Inhibitory roles of $Zn^{2+}$ on enzyme activity were tested via FRET-based enzyme assay[5]. Fluorogenic peptide substrate (Dabcyl)-KTSAVLQ↓SGFRKM-E (Edans)-NH2; (GL Biochem) contains the cleavage site of SARS-COV-2 Mpro (cleavage site represented by ↓). Cleavage of the peptide is marked with an increase in fluorescence from EDANS, which was monitored with microplate reader (Spectramax M3, Molecular devices) at 360 nm excitation and 460 nm emission wavelengths.

As DTT chelates Zinc ions, SARS-COV-2 Mpro was buffer exchanged into reaction buffer (20 mM Tris, 100 mM NaCl, 0.5 mM TCEP, pH 7.3) using PD SpinTrap G-25 column (GE) to remove DTT. 5 µL of Mpro at a final concentration of 200 nM was added to 35 µL pre pipetted reaction buffer in a black 96 well plate. For $IC_{50}$ calculation, 5 µL inhibitor at concentrations ranging from 25 µM to 12.2 nM (2-fold serial dilution), was added to the protein-containing reaction mixture and incubated at 25°C for 30 min with gentle shaking. The reaction was started by adding 5 µL substrate at a final concentration of 20 µM, immediately after which the relative fluorescence was read for 45 min. The total reaction volume was 50 µL. Data were normalized by considering negative control (protein heat inactivated at 60º C for 5 min) as 100% inhibition while treating positive control as 0% inhibition.

To test reversibility of $Zn^{2+}$ inhibition, 200 nM protein was first incubated with 500 nM zinc acetate for 30 min at 25º C with gentle shaking as described above. The reaction was then initiated with 20 µM substrate. 1 and 5 mM EDTA were added in separate wells at 28$^{th}$ minute when the reading was being taken.

**SARS-CoV-2 Mpro crystallization and soaking with Zinc:** Purified protein was diluted to 13.6 mg/ml for crystallization in buffer C. Several flower-like multi-crystals were obtained after overnight incubation at 20º C in the reservoir solution containing 100 mM Bis-Tris, 20% PEG 3350 and 5% DMSO pH 6.5[6]. These multi-crystals were used to prepare seeds using seed beads (Hampton research). Seeding was done into 3 µL protein: reservoir (2:1) drop in a 24 well sitting drop plate (Hampton research). Thereafter, single crystals with thin plate-like morphology were obtained after overnight incubation. Reservoir containing 10 mM Zinc glycinate or Zinc gluconate (TCI chemicals #G0215 and #G0277, respectively) was added to wells containing good quality crystals and soaked for 4 h. Crystals were then fished out and cryo-protected in a solution containing the reservoir with 20% glycerol. Subsequently, crystals were immediately flash-frozen into liquid nitrogen and stored for further X-ray diffraction and data collection. Multiple attempts to co-crystallize with zinc salts failed due to heavy precipitation of the protein. Also, soaking solutions containing $Zn^{2+}$ such as Zinc acetate or Zinc sulphate deteriorated the crystal quality.

**X-ray data collection, processing and refinement:** X-ray diffraction data for Zinc-soaked SARS-CoV-2 Mpro crystals were collected at XRD2 beamline[7], Elettra Sincrotrone Trieste at 0.99Å wavelength on a Dectris Pilatus 6M detector. Collected data were processed with autoPROC[8] and structure was determined by molecular replacement using Phaser-MR of Phenix crystallographic suite[9] using 6Y2F as search template. Initial model building was done with AutoBuild[10] module. Structure and map quality were further improved by manual building with Coot[11] and refinement with autoBUSTER[12]. Refinement statistics are summarised in Supplementary Table 2. Final model has $R_{work}$ and $R_{free}$ of 0.19 and 0.21 respectively. The structure has no Ramachandran outliers and 0.8 % side chain outliers. Figures were made with UCSF Chimera[13] and Maestro, Schrodinger suite[14] (Licenced to ICGEB).

**Molecular Dynamics:** Crystal structure of SARS-CoV-2 Mpro with Zinc (PDB: 7DK1) was prepared with protein preparation wizard of Schrodinger suite. Protonation states at pH 7.4±0.5 were created for the complex, explicit hydrogens were added to the structure, and zero bond order was created for $Zn^{2+}$ ion. Hydrogen bond optimization was done with ProtAssign and finally restrained minimization was performed using OPLS3e force field to obtain input structure for further calculations and analysis before performing Molecular Dynamic (MD) simulations.

To analyse the stability of the SARS-CoV-2 Mpro-Zinc complex, a 1 μs MD simulation was performed using Desmond[15] (Schrodinger) and the coordinates were saved at an interval of 50 ps. Simulation system was built using OPLS3e force field and solvated with TIP3P water model. Orthorhombic box with an edge length of 10 Å was set, ensuring a minimal distance between the atoms of protein complex and edge of the box. Counter ions were added to neutralize the system; further, 0.15 M NaCl was added to the solvated box as salt. The prepared systems were relaxed before the actual simulation by a series of energy minimization and short MD simulations, which mainly comprise of six relaxation steps while keeping the solute restrained. Briefly, in the first two steps, systems were relaxed with Brownian Dynamics NVT at T=10 K for 100 ps and 12 ps respectively. In step 3 and 4, NPT equilibration was done for 12 ps at 10 K with restrains on heavy solute atoms. At step 5, the pocket was solvated. Finally, in step 6 and 7 short NPT equilibrations were done for 12 and 24 ps respectively. The NPT ensemble was employed for the simulations with Nose-Hover chain thermostat and the Maryna-Tobias-Klein barostat. RESPA integrate was used with a time step of 2 fs. For short range of coulombic interactions, a 9 Å cut off was considered. Analysis of the simulation was done with simulation event analysis, Desmond.

**Cell culture and virus strain**: Vero E6 cells (African green monkey kidney cells) were purchased from ATCC, grown and maintained in Minimal Essential Media (MEM; HIMEDIA; AL047S) supplemented with 10 % FBS (HIMEDIA, RM10681), 2 mM L-Glutamine (HIMEDIA; TCL012), 100 U/ml penicillin, and 10 mg/ml streptomycin, in a 5 % carbon dioxide incubator with controlled humidity at 37 °C. For antiviral studies, SARS-CoV-2 strain, USA-WA1/2020 was used. All the virus infection and subsequent experiments using virus were performed in BSL-3 (virology) facility at ICGEB, New Delhi.

**Cell viability assay:** Cytotoxicity of Zinc acetate, Zinc glycinate, and Zinc gluconate on the viability and proliferation of the Vero E6 cells was evaluated using MTT assay. Cells were seeded at a density of 7000 cells per well in a 96-well plate. After allowing the cells to attach overnight, they were treated with varying concentrations of the above compounds. Treatment was done in MEM supplemented with 2% FBS for 48 h, at the end of which MTT assay was performed as per manufacturer's protocol. GraphPad Prism software was used to determine the $IC_{50}$ (50% inhibitory concentration). The absorbance (A) was measured at 570 nm and the percentage cell viability was calculated using the following formula:

Percentage cell viability = ($A_{570}$ of treated)/$A_{570}$ of Untreated) *100

**Anti-SARS-CoV-2 assay:** Anti-viral assays with Zinc acetate, Zinc gluconate, Zinc glycinate and Zinc acetate: quercetin mixture (1:2 molar ratio) were performed using a standard assay reported for SARS-CoV-2 and other viruses[16,17]. Vero E6 cells were seeded in 24-well plates, a day prior to infection. The following day, Zinc and other compounds were added to these seeded cells at maximum non-toxic concentration (100 μM, 70 μM and 100 μM respectively) followed by infection with SARS-COV-2 (Multiplicity of infection; MOI= 0.1). The treated and virus infected cells were incubated for 48 h (37 °C, 5% $CO_2$) following which the supernatants were harvested for viral quantification by plaque assay and qRT-PCR.

**Plaque assay:** For viral quantification, Vero E6 cells were seeded in 96 well plates, followed by viral inoculation on the next day using dilutions; starting at 1:50 the virus was double diluted till 1:51200. The virus was incubated with the cells for 2 h at 37 °C for viral adsorption. Thereafter, the media containing the inoculum was removed, and wells were overlaid with 150 μL of 1% carboxymethylcellulose (CMC) prepared in MEM media (containing 10% FBS). The plates were then incubated at 37 °C for 96 h with 5% $CO_2$ and 75% humidity. Post incubation, the cells were fixed with 5% formaldehyde before washing twice with 1× PBS and staining was performed using 0.25% crystal violet (prepared in 30% methanol). Plaques were visualized and counted to calculate viral titers using the following formula:

Plaque forming units (pfu) = (No. of plaques)/ (Dilution × volume of virus)

**qRT-PCR:** To quantify the viral RNA using qRT-PCR, 150 μL media from the treated, untreated and virus infected wells was collected, and used for RNA isolation using the NucleoSpin Viral RNA isolation kit (740956.250). Isolated RNA samples were then subjected to One-step qRT-PCR using QuantiTect qRT-PCR kit (Qiagen #1054498) and PIKOREAL 96 Real-Time PCR system (Thermo scientific). Data analysis was performed using a standard curve to calculate genome equivalents of SARS-CoV-2 in all the samples.
